# Tissue correction strategies for ^1^H-MRS and their effects on age-dependent metabolite alterations

**DOI:** 10.64898/2026.11.20.767915

**Authors:** Karen L Galindo, Elena I Ivleva, Kimberly L. Chan

## Abstract

**Purpose:** Proton magnetic resonance spectroscopy (1H-MRS) metabolite concentrations are confounded by partial volume effects, since gray matter (GM), white matter (WM), and cerebrospinal fluid (CSF) each contribute differently to the measured signal. The standard CSF-only correction assumes a uniform metabolite distribution across GM and WM, but prior work has shown that metabolite levels differ between tissues which can introduce an additional confound. As such, a single alpha correction addressing this GM/WM difference has been previously introduced for GABA+. This approach, however, assumes a fixed WM-to-GM concentration ratio regardless of age, sex, and other biological factors known to influence brain metabolism. Thus, this study introduces two more granular tissue correction approaches, a decade-pooled α and an individualized α, and compares them to the existing CSF-only and metabolite-pooled α corrections to evaluate how the choice of tissue correction approach affects the age-dependent slopes of glutamate + glutamine (Glx), myo-inositol (mI), total choline (tCho), total creatine (tCr), and total N-acetylaspartate (tNAA).

**Methods:** Data came from a public dataset of 102 healthy adults (49M/53F, ages 20–69) scanned at 3T with PRESS-localized MRS in the WM-dominant centrum semiovale (CSO) and GM-dominant posterior cingulate cortex (PCC). Metabolite concentrations were quantified using each of the four tissue correction methods: a CSF-only correction, and three α-based corrections (metabolite-pooled α, decade-pooled α, and individualized α), where α (ratio of pure WM to pure GM concentration) was derived by linearly regressing the CSF-only concentrations against GM fraction for both CSO and PCC across subjects grouped by all participants, by decade of age, or by individual, respectively. Concentrations were modeled as nonlinear functions of age (Generalized Additive Models) and linear (robust linear regression).

**Results:** The choice of tissue correction significantly altered metabolite age-relationships. mI, tNAA, Glx, and tCho age-dependent slopes were most sensitive to the choice of tissue correction method, while tCr was the least sensitive. Notably, both the decade-pooled α and individualized α corrections revealed nonlinear increases in mI and tCho with age that were not apparent using CSF-only or metabolite-pooled α correction. Sex-specific age effects were found for tCho and tCr, with increases primarily driven by female participants, potentially related to menopausal hormonal changes.

**Conclusion:** Tissue correction choice can influence the results of brain metabolism aging studies. As such, the choice of tissue correction method must be selected carefully depending on the design of 1H-MRS study, informed by whether a study is cross-sectional or longitudinal and whether bulk tissues changes are expected in the population of interest.

## Introduction

Proton magnetic Resonance Spectroscopy (^1^H-MRS) is a non-invasive technique that allows for *in vivo* quantification of brain metabolites. Since MRS data needs to be acquired from relatively large, predefined voxels to obtain sufficient sensitivity, the resulting metabolite signals are greatly influenced by the voxel’s tissue composition of gray matter (GM), white matter (WM), and cerebrospinal fluid (CSF) which each have different compound concentrations and magnetic properties. As normal aging^1^ and neuropsychiatric disorders such as Alzheimer’s disease,^2^ multiple sclerosis,^3^ and Huntington’s disease^4^ are often accompanied by significant alterations in tissue composition, these partial volume effects can introduce significant confounds into the metabolite measures. To account for these confounds when estimating metabolite concentrations, tissue correction methods are applied to adjust for differences in the water reference concentration and water T1 and T2 relaxation effects between tissues ^5–7^. The most common approach, the CSF-only correction, assumes that the measured signal originates only from the combined GM and WM volume and not the CSF^5,8^. This approach, however, only partially accounts for bulk tissue changes as it assumes that metabolite levels are equally distributed across GM and WM. In contrast, prior work has shown that metabolite levels are not equally distributed across these two tissue compartments with higher GABA+ (GABA + macromolecules) in GM than WM, for instance^5,9,10^. Considering the limitations of this standard approach, a tissue correction approach to account for partial volume effects in the metabolite levels themselves was previously introduced^5^. This correction was done by estimating an “alpha (α)” term, based on the GABA+ concentration ratio between WM and GM in a cohort of healthy adults^5^. This term was calculated from the GABA concentrations measured from five different voxel locations pooled across all participants and applied as a fixed correction factor to every participant’s data. Although this method was initially introduced for GABA+ only, it can be applied to other metabolites that have significant GM/WM differences.

A limitation to this approach, however, is that it assumes the metabolite concentration ratio between WM and GM is constant regardless of age and other biological factors that have been previously shown to affect brain metabolism^11–14^. This is especially pertinent in aging studies, as GM and WM are primarily governed by different metabolic processes and thus show different vulnerabilities to age-related processes such as demyelination in WM and microstructural atrophy in GM^1^. These differences come from their distinct tissue composition; WM is composed largely of lipids that form the myelin sheath^15^, while GM relies more heavily on glucose metabolism to support its higher signaling activity^16,17^. Furthermore, healthy aging across the adult lifespan has been shown to be associated with progressive declines in GM volume beginning in early adulthood, while WM volume increases until midlife before gradually declining^1^. As such, age-related changes in metabolite levels could potentially reflect this shift in tissue composition instead of true changes in neurochemistry unless properly accounted for. For instance, prior GABA-edited MRS studies have shown that age-based between-group differences that appear significant in uncorrected data can disappear or are substantially reduced depending on the choice of tissue correction method^5,8,18^. Besides age, brain metabolism in WM relative to GM could potentially differ between individuals as a result of sex, menopausal status, and genetic factors such as APOE genotype, BDNF Val66Met polymorphism and variants in the glutamate transporter gene SLC1A2, all of which have been linked to differences in tissue-specific metabolism^11,14,19–23^. Together, these variations in brain metabolism suggest that applying a single α value may not sufficiently account for inter-individual differences in tissue composition observed between individuals.

Considering the limitations of a one-size-fits-all approach to tissue correction, a more targeted correction such as a decade- or individualized-based α value could be applied to better account for the range of biological confounds that could affect brain metabolism. In these approaches, the original tissue correction framework is extended to incorporate different assumptions about how much the WM-to-GM metabolite relationship is shared between individuals; a decade-based α correction assumes that participants within the same decade of age share a similar metabolite distribution across WM and GM while an individual α correction value assumes no such relationship exists and needs to be estimated from the data on a case-by-case basis.

The goal of these targeted corrections is to more accurately characterize brain metabolite levels across the healthy adult lifespan, which is crucial for establishing a baseline for detecting pathological aging and informing targeted interventions to restore healthy brain metabolism. To date, the majority of MRS studies in heathy aging have relied on linear models across broad adult age ranges^12,24–26^. Linear models, however, cannot capture age-related changes that reverse direction or plateau over time, as these trajectories can be influenced by biological transitions associated with aging, e.g., hormonal fluctuations during menopause or declining brain energy metabolism during a midlife “critical window”^11,27^. Nonlinear models may better capture these patterns, but existing work on the lifespan trajectories of *in vivo* brain metabolism is limited; one prior study modeled nonlinear trajectories of specific metabolites, but only in participants ages 7–34, with middle and older adulthood remaining unexplored^28^. A second study modeled nonlinear trajectories across nearly the full lifespan but focused exclusively on GABA+, creating unanswered questions as to how other key metabolites change with healthy aging^29^. Beyond age, sex is another important factor to consider separately, as sex-specific metabolite changes have been observed with multiple metabolites^26,30^. As such, the goal of this study is to evaluate different α tissue correction methods for ¹H-MRS, and to examine how different correction approaches affect non-linear age-dependent effects (cross-sectional “trajectories”) of metabolite concentrations as a function of sex across the adult lifespan.

## Methods

### In Vivo Data

All the data used for this study originate from a publicly available dataset of 102 healthy adults (49 M/53 F, ages 20– 69) with approximately equal numbers of men and women in each decade of life^2626,31^. This data was acquired on a clinical 3 Tesla MRI scanner (Ingenia CX, Philips Healthcare, The Netherlands) in two 30 × 26 × 26 mm^3^ voxel locations: the centrum semiovale (CSO, predominantly WM) and posterior cingulate cortex (PCC, predominantly GM). Data for each participant included a T1-weighted MPRAGE scan acquired at 1 mm^3^ isotropic resolution and water-suppressed metabolite spectra acquired with PRESS localization, VAPOR water suppression^32^, an echo time (TE) of 30ms, a repetition time of 2000 ms and 96 transients. A metabolite-nulled macromolecule (MM) spectrum was also acquired for each participant from the same voxels using the same acquisition parameters as the water-suppressed metabolite data, except with CHESS water suppression^33^ and a 600 ms inversion time^31^. An unsuppressed water reference was also acquired for each participant from the same voxels. As previously reported, data was of high-quality for both regions with high mean N-acetylaspartate (NAA) signal-to-noise-ratios (SNR) of 160 for the CSO and 150 for the PCC, and narrow mean NAA linewidths of 6.8 Hz for the CSO and 7.0 Hz for the PCC^26,34^. In the PCC voxels of this data set, the GM fraction was shown to decrease moderately with age while the WM fraction increased weakly with age, indicating significant changes in bulk tissue composition with age that need accounting for^31^. Table S1 provides the minimum reporting standards for *in vivo* MRS^35^.

### Data Processing

Data preprocessing was performed in Gannet 3.1^36^. Residual water was removed from the time-domain signal using a Hankel singular value decomposition (HSVD) filter^37^. Spectra were zero-filled to 4096 points and apodized with 1 Hz exponential line broadening. Spectra were manually zero-order phase corrected and frequency-aligned.

All spectral fitting was performed in ProFit-1D^38^. Fit quality was assessed using the fit quality number (FQN) calculated for each spectrum as previously described^38^. The FQN refers to the ratio of residual variance to noise variance, with lower values indicating a better fit. Metabolite basis sets were simulated in FID-A^39^ using previously-published J-coupling constants and chemical shifts^40,41^. These simulations were performed at a TE of 30ms over a 36 × 36 mm^2^ spatial plane corresponding to the two refocusing-pulse directions, which was split into 19 x 19 segments, and apodized with 3 Hz exponential line broadening. The basis sets for both regions included glutamate (Glu), glutamine (Gln), NAA, creatine (Cr), phosphocreatine (PCr), glycerophosphocholine (GPC), phosphocholine (PCh), myo-inositol (mI), taurine (Tau), glutathione (GSH), N-acetylaspartylglutamate (NAAG), aspartate (Asp), scyllo-inositol (Scyllo), lactate (Lac), γ-aminobutyric acid (GABA), glucose (Glc), and glycine (Gly). A clean MM spectrum for each region, which was averaged over all participants for each region separately, was also obtained by fitting the inversion-recovery spectra with a limited metabolite basis set containing total creatine (tCr; Cr, PCr), total choline (tCho; GPC, PCh), total NAA (NAA, NAAG), and Glx (Glu, Gln) and extracting the spline baseline containing only the MM as previously described^31^. The resulting CSO and PCC MM spectra were included as an additional basis component alongside the simulated metabolite bases to fit data from their respective regions. Tissue segmentation (GM, WM, CSF fractions) was performed in SPM12^42,43^ as implemented in Gannet 3.1^36^. The resulting tissue fractions, denoted as *f_GM_*, *f_WM_*, and *f_CSF_* represent the proportions of gray matter (GM), white matter (WM), and cerebrospinal fluid (CSF), respectively, within each MRS voxel. Spectra from 14 subjects were excluded from the CSO and PCC analyses: 12 (11.7%) due to segmentation errors and 2 (1.96%) due to noisy spectra.

### Tissue Correction Methods

Four tissue correction approaches were applied and compared: CSF-only correction, metabolite-pooled α correction, decade-pooled α correction, and individualized α correction.

Absolute metabolite concentrations for mI, Glx, tCho, tCr, and tNAA relative to unsuppressed water were calculated using the widely-used Gasparovic method as previously described^6^. These metabolites were selected because they can be robustly and reliably quantified from a short-TE PRESS spectrum without the need for specialized acquisition methods. For this study, GM and WM water T1 and T2 relaxation times were taken from Wansapura et al.,^44^and CSF water T1 and T2 were taken from Lu et al.^45^ and Piechnik et al.,^46^ respectively. Metabolite-specific T1 and T2 relaxation times were taken from previously reported values for Glu, Gln, Ins, Cr, PCr, NAA, NAAG, GPC, and PCh.^47–49^ Water concentrations for each tissue compartment were calculated relative to a tissue water density of 55.51 mol/kg from Gasparovic et al.^6^ based on values originally reported by Ernst et al.^50^ (fGM = 0.78, fWM = 0.65, fCSF = 0.97), yielding compartmental water concentrations of 43.30 mol/kg in GM, 36.08 mol/kg in WM, and 53.84 mol/kg in CSF.

The first of four tissue correction methods, CSF-only correction, assumes the ¹H-MRS signal originates only from GM and WM^6,50^, an assumption that holds for the metabolites-of-interest for this study^51^. Under this assumption, the measured concentration is expressed as:

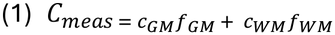

where *C_GM_* and *C_WM_* represent the concentration of each metabolite in GM and WM respectively, with CSF excluded. *C_meas_* is then normalized by the combined GM and WM signal:

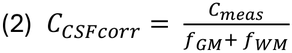

This approach, however, assumes uniform metabolite concentrations across GM and WM and does not account for potential differences in metabolite distribution between the two tissue types. For example, tCho has been found to be present in higher concentrations in WM compared to GM while the opposite has been found to be true for Glx^52^.

As such, a second correction method that accounts for these potential differences was also explored. Introduced previously by Harris et al.^5^, this method builds upon the CSF-only correction in equation 2 with the inclusion of an additional α correction term in the denominator that accounts for the relative metabolite concentration between GM and WM:

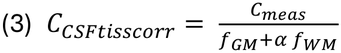

where α is the expected metabolite concentration ratio in WM relative to GM:

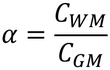

Setting α = 1 indicates no difference in metabolite concentration between WM and GM and thus reduces this equation to the CSF-only corrected approach above; setting = 0 assumes a purely GM voxel. Since α values are defined relative to the GM concentration, metabolites present at higher concentration in WM than GM, such as tCho, will have an α value greater than 1, while metabolites present at higher concentration in GM than WM will have an α value less than 1. Initially demonstrated for GABA+, the original α was set to 0.5 to account for the two-times higher GABA+ in GM than in WM but can be experimentally derived for other metabolites. This was done here across data acquired from all 88 participants which were acquired in two regions with significantly different tissue compositions (i.e. the WM-heavy CSO and the GM-heavy PCC), thus allowing for an estimation of α using a linear regression analysis.

This was done by plotting the CSF-only corrected concentration against GM fraction (*f_GM_*), defined as 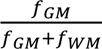 and performing a linear fit using “robust fit”, an iteratively reweighted least squares with a Tukey’s biweight function^53^ to reduce the influence of outliers:

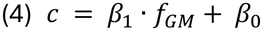

where *β*_1_ is the slope and *β*_0_ is the intercept. *β*_0_ represents the metabolite concentration if the voxel consisted of purely WM (*f_GM_* = 0, *C_WM_*). Extrapolating the same line with *f_GM_*= 1 yields the concentration for a purely GM voxel (*C_GM_*= *β*_0_ + *β*_1_). From this fit, α can then calculated as the ratio between these two extrapolated concentrations:

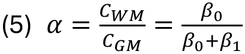

Unlike Harris et al.^5^, we did not perform a subsequent group normalization step in which subject-level concentrations are rescaled to an average voxel composition. Instead, concentrations were expressed on the pure-tissue scale instead of the average-voxel-normalized scale. This was done to keep the metabolite measures independent of the cohort composition, since bulk tissue proportions varied substantially across subjects due to the wide age range included in this cohort.

Since the metabolite-pooled α approach assumes a static value across individuals of all ages which may not properly reflect the biological variation present in healthy adults, we also calculated decade-pooled α values and individualized α values. Both follow the same calculation as metabolite-pooled α but differ in subject grouping before fitting. For the decade-pooled α, subjects were grouped by decade of life (20s, 30s, 40s, 50s, and 60s) and a robust linear regression^53^ was performed over the CSF-only corrected concentrations and GM fractions within each group, yielding one α per metabolite per decade. For individualized α, a linear model was fit across two voxels of differing tissue composition in each participant which yielded a participant-specific α per metabolite. Participants with negative α values, which are physiologically implausible, were excluded (2 subjects (1.96%), for mI only) as these values likely reflect unreliable concentration estimates.

### Extracting Age-Dependent Effects

To evaluate the effect of the different tissue correction methods on the age-dependent slopes of Glx, mI, tCho, tCr, tNAA, both linear and nonlinear models were fit to the age-based metabolite data and evaluated for curve differences. For linear regressions we used “robust fit” in MATLAB using a Tukey’s biweight function^53^. Nonlinear models were carried out using Generalized Additive Models (GAMs) in R (version 4.6.0) using the mgcv package^54–58^ ^59^. Unlike traditional linear regression methods, GAM does not require the relationship between predictor and outcome to be specified beforehand, and the shape of the relationship is estimated directly from the data using smooth nonparametric functions^60^. This allows GAMs to capture nonlinear patterns including inflection points that may not be captured by an isolated linear or polynomial fit without overfitting^59^. Age-associated effects, as estimated with GAMs, were modeled as a smooth term using thin-plate regression splines, the default basis function in mgcv^54,57,59^. Smoothing parameters were estimated using restricted maximum likelihood (REML), which penalizes excessive curvature^54^. The basis dimension was set to k=10 for all smooth terms; this choice was verified using the k-index diagnostic (gam.check) in mgcv^54,60^. Each metabolite correction method was compared to every other method using an ordered-factor GAM, yielding six pairwise smooths comparisons. Method-specific age smooths were fit for each metabolite and voxel. A random intercept for subject was included, since each subject contributed a paired measurement under every method.^55^ To assess sex-based differences in the metabolite slopes with age, separate GAMs were fit to male and female participants. Differences in age-based effects between sexes were tested by modeling sex as an ordered factor, with a shared smooth and a difference smooth capturing sex-specific deviation^55^.

### Statistical Analysis

Pairwise differences in individualized α values were assessed using the Wilcoxon rank-sum test^61^. Pairwise differences in metabolite concentrations corrected with an individualized α across participants split into younger (age 20–39 years) and older (age 50–69 years) female and male groups was also assessed with a Wilcoxon rank-sum test. This assessment was performed to provide greater power in detecting sex-by-age differences as splitting the data further by sex reduces the overall statistical power. GAMs diagnostics and model outputs were assessed using standard diagnostic functions in mgcv^54–58^. The shape of each fitted smooth term was summarized using its effective degrees of freedom (edf), which reflects the estimated linearity of the metabolite-age relationship: an edf close to 1 indicates a linear relationship, an edf close to 2 indicates a quadratic-like curve, and an edf greater than 2 indicates a progressively more complex, nonlinear pattern^55^. Robust regression uses a t-test on a single, fixed slope. GAMs significance testing works differently because the shape of the curve itself is estimated from the data using REML, a method that estimates how much smoothing to apply by mathematically separating out the influence of the model’s fixed effects^54^. The p-value for each smooth term is then estimated by mgcv’s summary.gam() function, which uses a chi-squared-type test based on the term’s estimated coefficients, their covariance, and its edf, instead of a single slope. The F-statistic and reference degrees of freedom were used to assess the significance of the smooth terms. To identify where two smooths diverged, the difference between fitted curves at each age point was calculated, along with its 95% confidence interval; age ranges where this interval excluded zero were considered significantly divergent. For sex-comparison models this was computed using the tidygam package^60,62^; for pairwise method-comparison models with a subject-level random intercept, it was computed directly from the model’s linear predictor matrix and coefficient covariance matrix. For comparisons involving multiple simultaneous tests, p-values were corrected for multiple comparisons using the false discovery rate (FDR) method^63^. P-values were considered statistically significant if p < 0.05. FDR-corrected p-values are reported throughout, with findings significant only before correction labeled as uncorrected.

## Results

Representative voxel placements and example spectral fits for the WM-dominant centrum semiovale (CSO) and GM-dominant posterior cingulate cortex (PCC) are shown in Figure 1a and Figure 1b, respectively. For both regions, spectral fits were of high quality, with close agreement between the overall spectral model (red) and the data (black); this was also reflected in low median FQN values of 11.3 (IQR: 8.7–15.2) in the CSO (n=88) and 10.7 (IQR: 7.9–13.9) in the PCC (n=88). As expected, the CSO and PCC voxels showed distinct tissue compositions, with mean GM fractions of 19.2% and 68.4%, respectively (Figure 1c).

**Figure 1.**
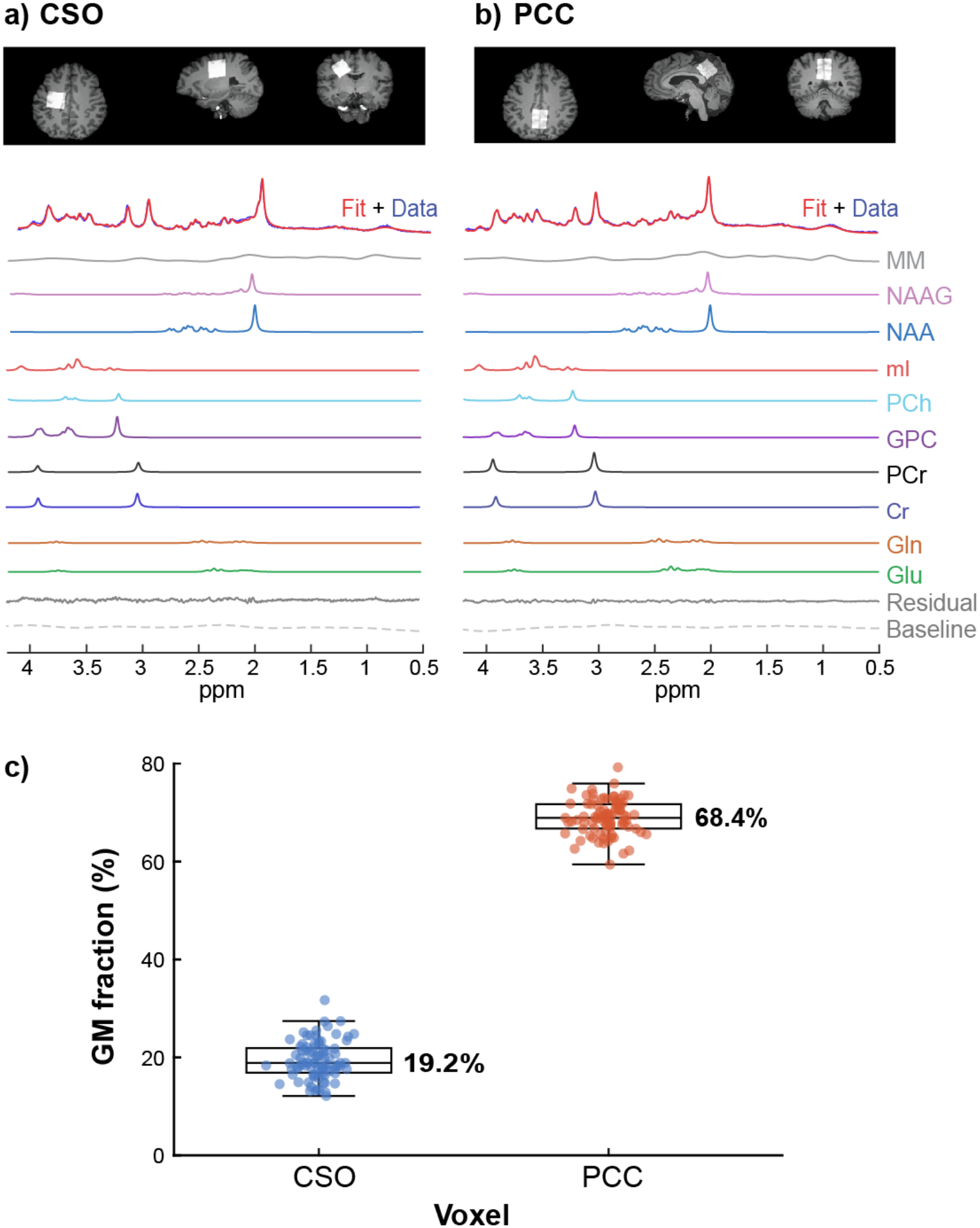
Voxel placement, representative spectral fits, and tissue composition. Representative voxel placement and ProFit-1D spectral fits were of high-quality for both the centrum semiovale (CSO, a) and posterior cingulate cortex (PCC, b) **(c)** Boxplots showing the gray matter (GM) fraction for each voxel across all participants. The mean GM fraction of 19.2% for the CSO (white matter–predominant) and 68.9% for PCC (gray matter– predominant).

Figure 2 shows representative linear regression fits to estimate α for each tissue correction method using Glx and tCho as examples. Across all three group levels, Glx and tCho showed opposing relationships with GM fraction, consistent with their known tissue specific distributions and was consistent across all three levels of grouping ^52^. Glx levels were positively related to GM fraction, yielding α values below 1 across all three tissue correction methods (α = 0.29–0.31; Figure 2a). In contrast, tCho levels were negatively associated with GM fraction, yielding α values above 1 across all three tissue correction methods (α = 1.24–1.38; Figure 2b).

**Figure 2.**
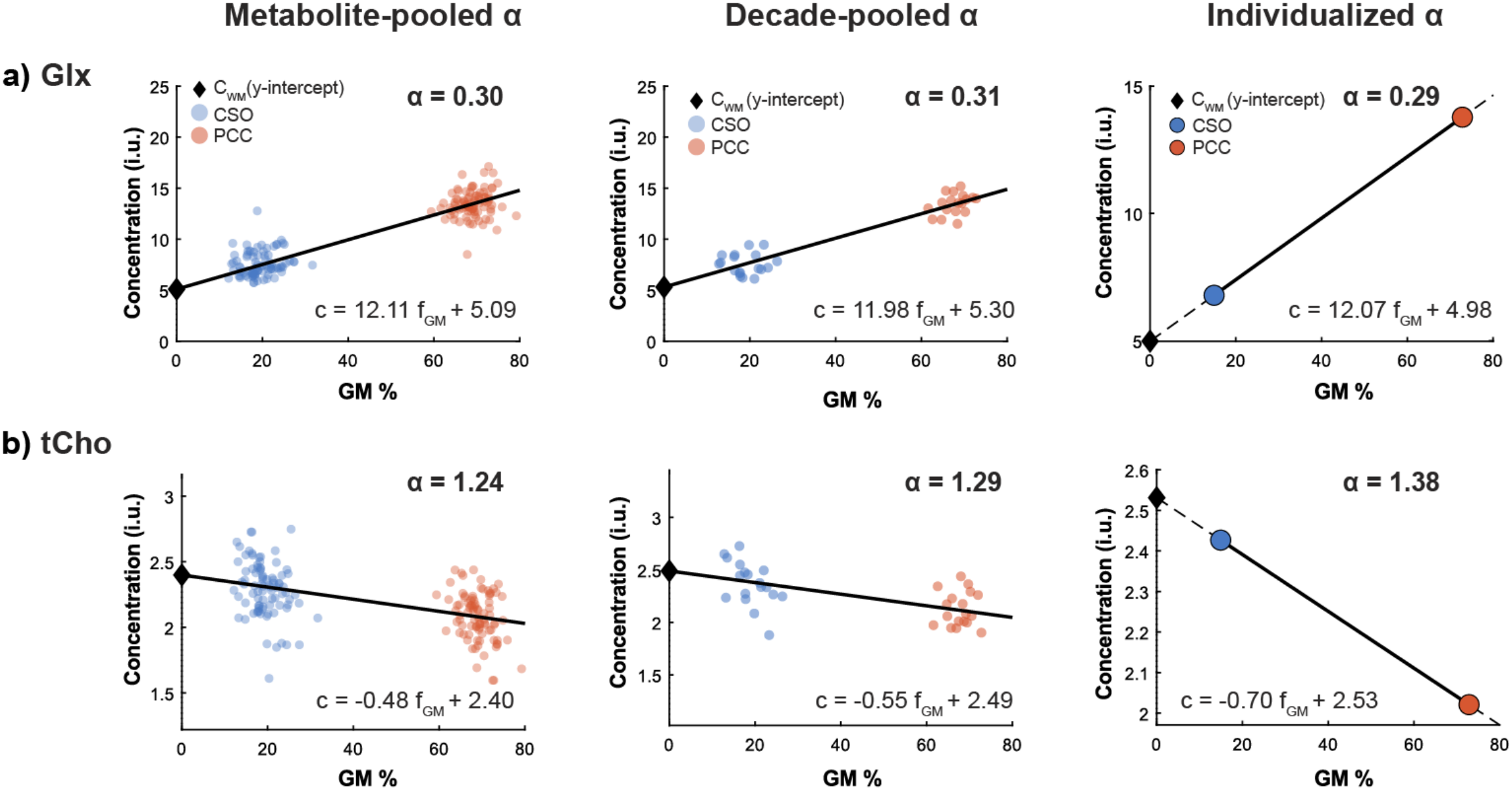
Calculation of metabolite-specific α values using individualized and group-level approaches. The metabolite-pooled α was calculated by performing a linear regression to the metabolite data taken from all participants in the cohort. The decade-pooled α was calculated by fitting a linear regression model to the metabolite data taken from all participants in the same decade of age. The individualized α was calculated for each subject by fitting a line between the CSF-only corrected metabolite concentrations from the CSO (blue) and PCC (orange) voxels as a function of GM fraction. Representative α estimations are shown for Glx (a) and tCho (b). Black diamonds denote the extrapolated pure WM concentration, and solid black lines indicate the linear regression model used to estimate α. Across all α estimation approaches, the α values for Glx are less than 1, indicating higher Glx levels in gray matter than in white matter, while the α values for tCho are greater than 1, indicating lower tCho levels in gray matter than in white matter, consistent with prior literature.

### Individualized and pooled α values across metabolites and age

Figure 3a shows boxplots of individualized α values plotted alongside metabolite-pooled α values. Here, it can be seen that the α values differed substantially between all five metabolites (Figure 3a). Of these five metabolites, tCho demonstrated the highest α values, indicating greater levels in WM vs. GM, while Glx demonstrated the lowest α values, indicating greater levels in GM vs. WM. While the metabolite-pooled α values (Glx = 0.30, mI = 0.69, tCho = 1.24, tCr = 0.63, tNAA = 0.99) closely matched the median individualized α values for each metabolite (Glx = 0.28, mI = 0.71, tCho = 1.22, tCr = 0.61, tNAA = 1.17), the individualized α values still span a moderate range about the median.

**Figure 3.**
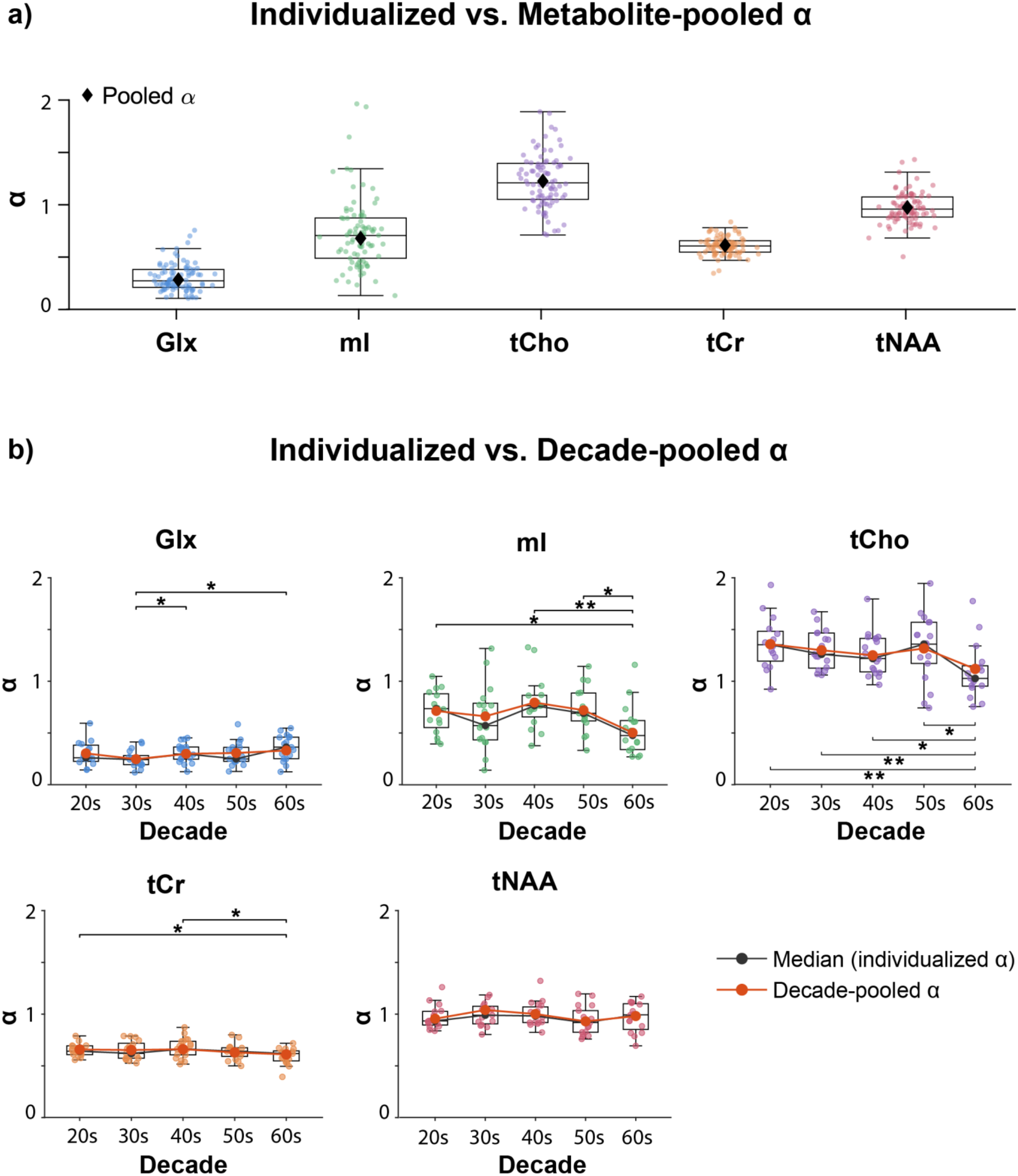
Comparison of individualized α values with metabolite-pooled α and decade-pooled α values across metabolites and age. **(a)** Boxplots showing the distribution of the individualized α values across metabolites, with corresponding metabolite-pooled α estimates (Glx = 0.30, mI = 0.69, tCho = 1.24, tCr = 0.63, tNAA = 0.99) for comparison. **(b)** Boxplots showing the distribution of the individualized α values across each decade of age, with corresponding decade-pooled α values for comparisons. Significant differences are indicated by brackets and asterisks (*p < 0.05, **p < 0.01). Statistically significant differences in α values across decades of age were found for Glx, mI tCho, and tCr.

Examining this variation more closely and grouping the individualized α values by decade (Figure 3b) revealed that the α values for most metabolites vary with age. Glx α values were found to increase with age, with significantly lower values in participants in their 30s relative to those in their 40s (p = 0.037) and 60s (p = 0.012). In contrast, mI α values showed significant decreases with age, with significantly lower α values in participants in their 60s relative to those in their 20s (p = 0.019), 40s (p = 0.005), and 50s (p = 0.011). Similarly, tCho α values showed a similar age dependency with significantly lower α values in participants in their 60s relative to those in their 20s (p = 0.003), 30s (p = 0.004), 40s (p = 0.010), and 50s (p = 0.018). tCr α values were also found to be significantly lower in participants in their 60s relative to those in their 20s (p = 0.05) and 40s (p = 0.023). For all five metabolites, the median individualized α per decade closely matched the corresponding decade-pooled α’s which are shown in Table 1. However, some variability between individuals can still be observed within each decade-of-age.

**Table 1:** Decade-pooled α values for each metabolite across five age decades.

| **Metabolite** | **20s** | **30s** | **40s** | **50s** | **60s** |
|---|---|---|---|---|---|
| **Glx** | 0.302 | 0.247 | 0.298 | 0.307 | 0.330 |
| **mI** | 0.728 | 0.676 | 0.812 | 0.736 | 0.514 |
| **tCho** | 1.326 | 1.268 | 1.220 | 1.286 | 1.092 |
| **tCr** | 0.640 | 0.635 | 0.642 | 0.612 | 0.593 |
| **tNAA** | 0.957 | 1.042 | 1.003 | 0.932 | 0.983 |

### Effect of tissue correction method on metabolite concentration cross-sectional “trajectories” across age

Figure 4 shows the age-based slopes for all five metabolites using each of the different tissue correction methods in the PCC.

**Figure 4.**
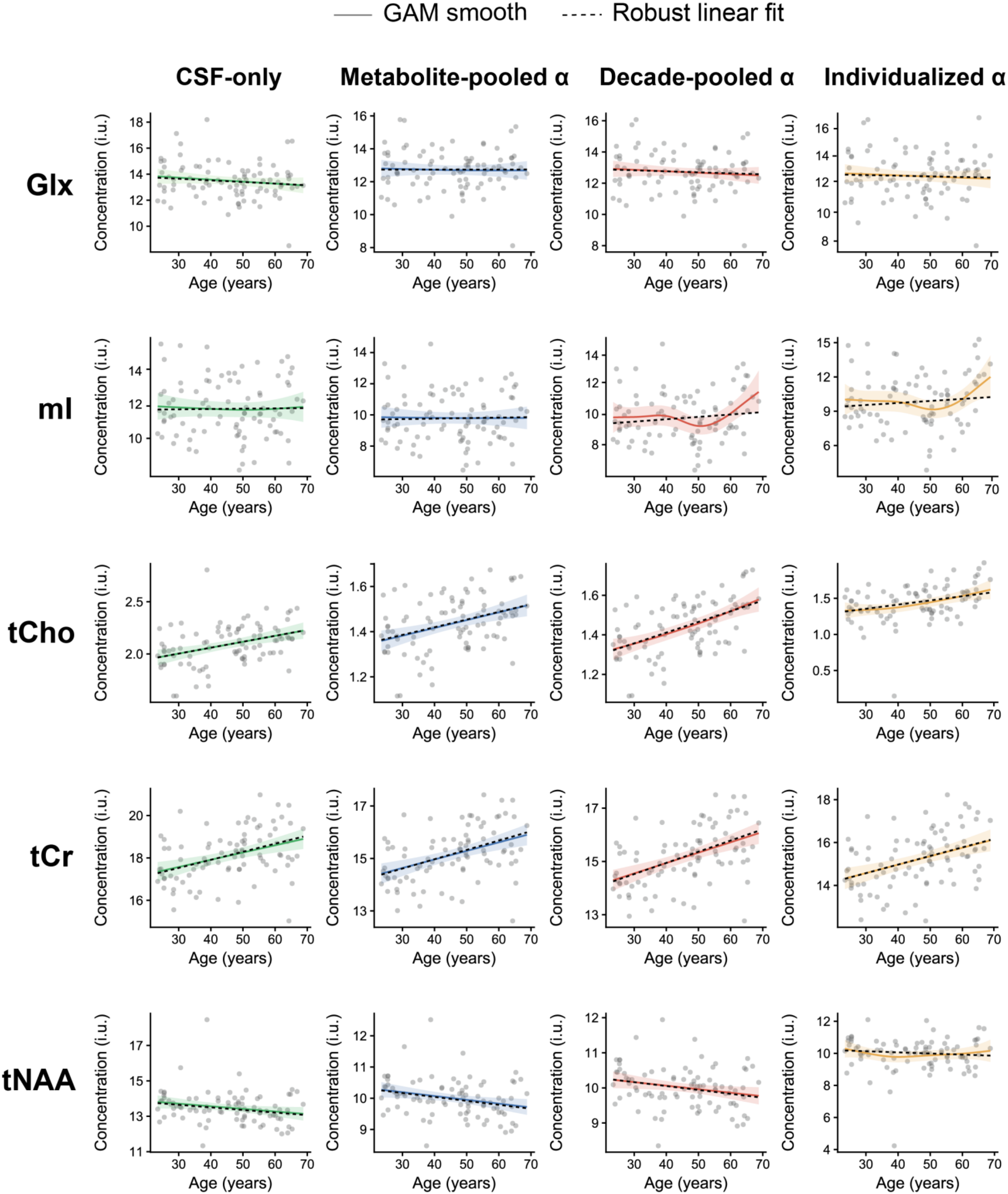
GAM-based smooths of metabolite concentrations in the PCC, by tissue correction method. Glx, mI, tCho, tCr, and tNAA concentrations and their corresponding GAM-based smooth functions are plotted against age for all four correction methods. Solid lines show the GAM smooth fit with shaded 95% CI; dashed lines show the robust linear fit. Individual subject values are overlaid as points. Significant metabolite-age effects with both GAMs and robust linear regression were found for tCho and tCr across all four correction methods (all p ≤ 0.0028), tNAA with the CSF-only, metabolite-pooled α, and decade-pooled α corrections (all p ≤ 0.043). A significant metabolite-age effect with GAMs was found for mI with the decade-pooled α correction (p = 0.049).

#### Glx

While Glx levels were found to decrease linearly with age (edf = 1.00) across all tissue correction methods, this was not found to be statistically significant with either GAMs (all uncorrected p > 0.18) or robust linear regression (p > 0.39) modeling. However, the Glx-age slope shape differed significantly between correction methods across the full age range of 24–69 years. The CSF-only trajectory (slope = -0.0088) diverged significantly from both the metabolite-pooled α trajectory (slope = +0.0039; p = 0.0012) and the decade-pooled α trajectory (slope = -0.0024; p = 0.045, uncorrected), and the metabolite-pooled α trajectory diverged from the decade-pooled trajectory as well (p < 0.0001) across the same age range.

#### mI

GAM modeling revealed a positive nonlinear increase in mI with age only when using a decade-pooled α correction (edf = 3.31, uncorrected p = 0.0496). No mI-age effect was found when using a CSF-only or a metabolite-pooled α correction, however, with either GAM or robust linear regression modeling (uncorrected p > 0.85). Nonlinear increases in mI with age were also found with GAMs with an individualized α correction (edf = 3.03); however, this result was not statistically significant (uncorrected p = 0.092). Notably, the shapes of mI-age slopes also differed between methods. The mI-age trajectory corrected for CSF-only (slope = +0.0003) diverged significantly from the mI-age trajectory corrected with a decade-pooled α (slope = +0.0154) across the full age range (24–69 yrs; p < 0.0001). The mI-age trajectory corrected for CSF-only also diverged from the mI-age trajectory corrected with an individualized α (slope = +0.0178) over the same age range (p = 0.0157, uncorrected). The mI-age trajectory corrected with an individualized α diverged from the mI-age trajectory corrected with a metabolite-pooled α (slope = +0.0019) in older age (62–69 yrs; uncorrected p = 0.0217). The mI-age trajectory corrected with decade-pooled α diverged from the trajectory corrected with metabolite-pooled α across the full age range (24–69 yrs; p < 0.0001).

#### tCho

Significant increases in tCho with age was observed for all four correction methods with both GAMs and robust regression (p ≤ 0.007). While linear tCho-age relationships were found when using a CSF-only correction and metabolite-pooled α correction (edf = 1.00), nonlinear increases in tCho with age were found when using the decade-pooled α (edf = 1.43) and individualized α (edf = 1.72) tissue correction approaches. The shape of the tCho-age trajectory also differed significantly between the CSF-only (slope = +0.0057) and metabolite-pooled α (slope = +0.0033) correction approaches across the full age range (24–69 yrs; p = 0.018), as well as between the metabolite-pooled α and decade-pooled α (slope = +0.0054) corrections over the same range (p < 0.0001)

#### tCr

A positive linear association (edf = 1.00) between tCr and age was found for all four correction methods with both GAMs and robust regression (all p ≤ 0.0008). The shape of the tCr-age trajectory also differed significantly between the CSF-only (slope = +0.0361) and decade-pooled α (slope = +0.043) correction approaches across the full age range (24–69 yrs, uncorrected p = 0.019), as well as between the metabolite-pooled α (slope = +0.036) and decade-pooled α corrections over the same age range (p < 0.0001).

#### tNAA

Linear decreases in tNAA levels with age were found with both GAMs and robust regression for the CSF-only, metabolite-pooled α, and decade-pooled α tissue correction approaches (uncorrected p < 0.05). Applying an individualized α correction, however, removes the tNAA-age relationship for both GAMs and robust linear regression (edf = 2.19, p = 0.44). Significant differences in tNAA-age trajectories across the full age range of 24-69 years were found between the CSF-only (slope = -0.0136) and decade-pooled α (slope = -0.0105) correction methods (uncorrected p = 0.040), as well as between the metabolite-pooled α (slope = -0.0120) and decade-pooled α corrections over the same age range (p < 0.0001).

### Effect of sex on age-related metabolite concentration cross-sectional “trajectories” across tissue correction methods

Metabolite age-based slopes fit separately to males and females for each metabolite and tissue correction method are shown in Figure 5.

**Figure 5.**
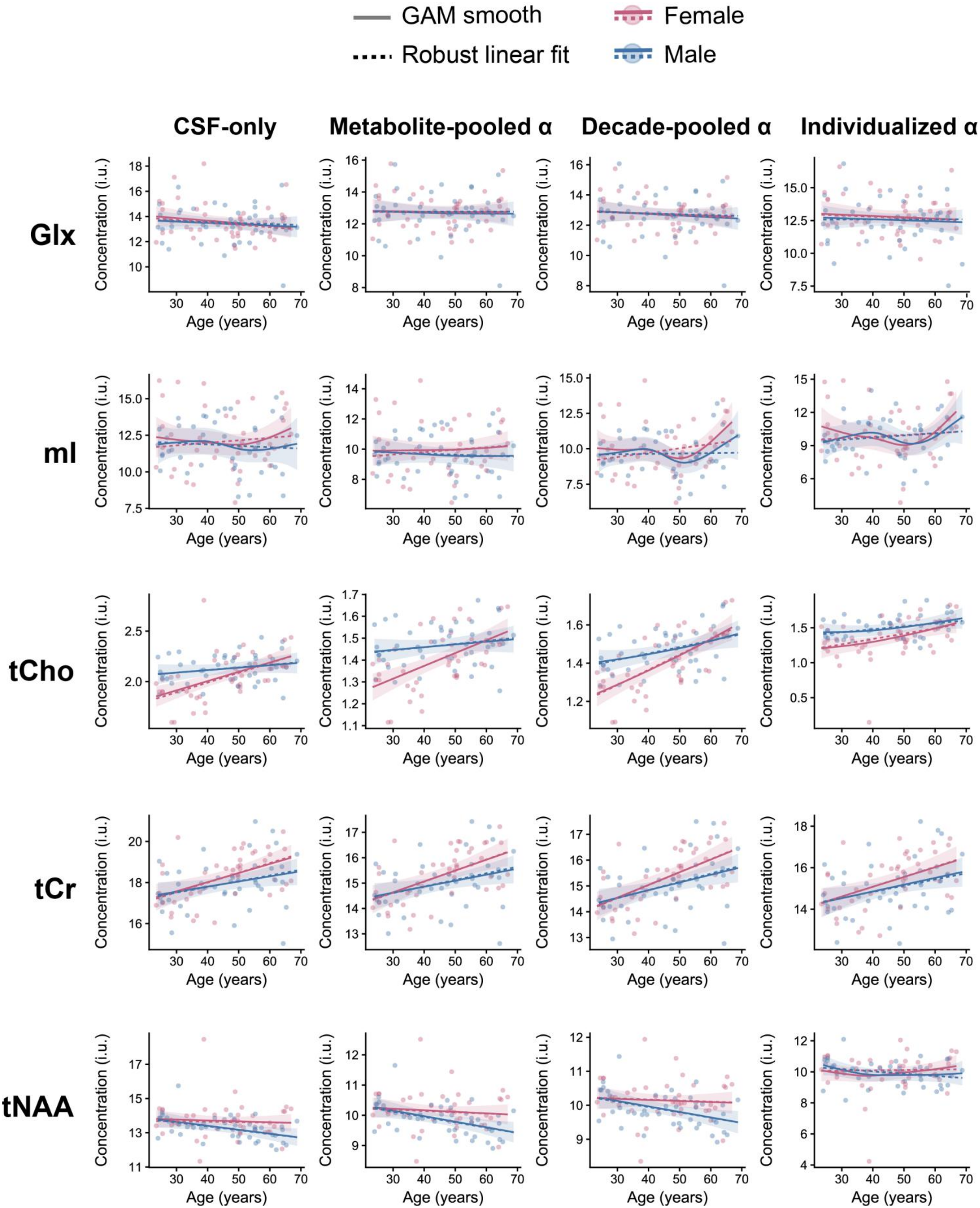
GAM-based smooths of metabolite concentrations, by sex and tissue correction method. Glx, mI, tCho, tCr, and tNAA concentration are plotted against age for females (pink) and males (blue), across all four correction methods. Solid lines show the GAMs smooth fit; dashed lines show the robust linear fit. Shaded regions indicate 95% CI; individual subject values are overlaid as points. Robust linear regression and GAMs showed female-specific effects for tCho and tCr across all four methods (p < 0.02) and male-specific effects for tNAA (CSF-only, metabolite-pooled α, decade-pooled α; all p ≤ 0.025), tCr with a decade-pooled α, metabolite-pooled α, and individualized α correction (p ≤ 0.039), and tCho with decade-pooled α and individualized α corrections (p ≤ 0.034).

#### Glx

No significant age effect was found for Glx in either females (all edf = 1.57-2.33) or males (all edf = 1.00) with any tissue correction method as modeled by GAM and robust regression (p > 0.2). Additionally, no significant trajectory differences were found between males and females for any method (p = 0.69 – 0.92).

#### mI

Similar to Glx, no significant age effect was found for mI in either females or males across all correction methods as determined by both GAM and robust regression (p > 0.09; edf, female/male: CSF-only 1.90/1.00, metabolite-pooled α 1.79/1.00, decade-pooled α 2.44/2.88, individualized α 2.38/2.77). Additionally, no significant trajectory differences were found between males and females for any method (p = 0.42 – 0.60).

#### tCho

A significant age effect was found for tCho in females across CSF-only (edf = 1.00), metabolite-pooled α (edf = 1.00), and decade-pooled α (edf = 1.14) correction approaches (all p < 0.01). This significant tCho-age effect was also found when using the individualized α correction approach (edf = 2.11; uncorrected p = 0.022). In males, significant age effects for tCho were found only when using the decade-pooled α (edf = 1.00; p = 0.034) and individualized α (edf = 1.00; uncorrected p = 0.020) but not the CSF-only (edf = 1.00; p = 0.13) and metabolite-pooled α approaches (edf = 1.00; p = 0.25). Trajectory differences between males and females from ages 24–48 were also observed for the CSF-only (slope: female = +0.0096, male = +0.0026), metabolite-pooled α (slope: female = +0.0057, male = +0.0014), and decade-pooled α (slope: female = +0.0078, male = +0.0033) correction methods with both GAM (uncorrected p < 0.05) and robust linear regression (p < 0.05). Although there is an apparent increase in tCho in both sexes, females exhibited lower levels until around age 48, followed by a steeper increase that narrowed the gap with males.

#### tCr

In females, tCr showed significant non-linear increases with age across all four tissue correction methods with both GAMs (edf: CSF-only - 2.74, metabolite-pooled α - 2.99, decade-pooled α - 2.91, individualized α - 2.61) and robust linear regression (p = 0.0007-0.011). In males, a significant tCr-age effects with GAMs were found only when performing decade-pooled α (edf = 1.00; p = 0.045), metabolite-pooled α (edf = 1.00; uncorrected p = 0.039) and individualized α (edf = 1.00; uncorrected p = 0.024) corrections and are consistent with what was found with robust regression (p ≤ 0.0191). No significant tCr-age effects were found in males when using a CSF-only correction (edf = 1.00 p = 0.063). No significant differences in trajectory shape were found between males and females for any tissue correction method (uncorrected p > 0.21).

#### tNAA

No significant tNAA-age effect was found in females across all tissue correction methods (edf = 1.00 for CSF-only, metabolite-pooled, and decade-pooled; edf = 2.21 for individualized), with either GAMs (p = 0.56–0.68) or robust regression (p = 0.54–0.95). In males, a significant tNAA-age effect was found with both GAMs (all edf = 1.00) and robust linear regression for the CSF-only, metabolite-pooled α, and decade-pooled α correction methods (p < 0.05) but not the individualized α correction method (p > 0.11). No significant differences in trajectory shape were found between males and females for any of the tissue correction methods (p = 0.15–0.23).

### Sex differences in individualized α derived concentrations across metabolites and age

Figure 6 demonstrates metabolite levels when corrected with individualized α values in younger (20–39 years) and older (50–69 years) males and females. tCho concentrations were significantly higher in younger males than in younger females (p = 0.003), and significantly lower in younger females than in older females (p = 0.006). tCr levels were significantly lower in younger females than in older females (p = 0.001).

**Figure 6.**
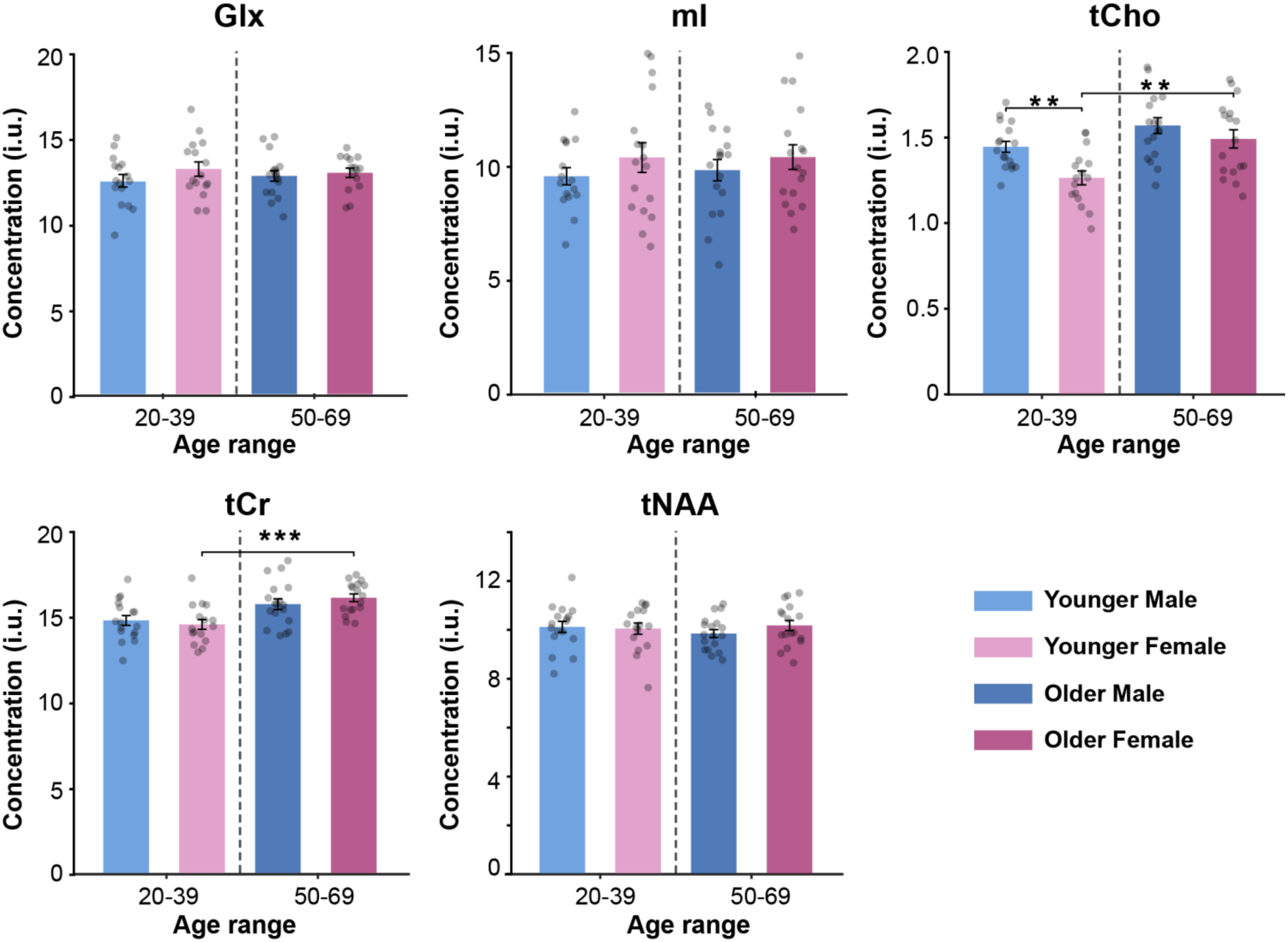
Individualized α corrected metabolite concentrations in the PCC by sex and age group. Concentrations for Glx, mI, tCho, tCr, and tNAA are shown separately for younger (20–39 years) and older (50–69 years) males and females. Bars represent mean ± SEM, and black dots represent individual subjects. tCho concentration was significantly higher in younger males than younger females (p < 0.003), and significantly lower in younger females than older females (p = 0.006). tCr concentration was significantly lower in younger females than older females (p = 0.001). No significant group differences were observed for Glx, mI, or tNAA. P values are indicated as *p* < 0.05 (*), *p* < 0.01 (**), and *p < 0.001* (***).

## Discussion

Biological and environmental variations such as genetics, hormonal effects, and lifestyle factors as well as technical differences in ROI definitions, spectral fitting and processing^64^ can all influence MRS outcomes; therefore, accounting for sources of variability such as tissue composition, becomes especially important. The goal of our study was to emphasize a neurobiological parameter that is often overlooked in MRS analyses: metabolites are not present in equal concentrations in gray and white matter, which presents a significant confound for studies where changes in bulk tissue composition are expected, such as studies of aging. In this study, we introduced two more granular α-based tissue correction approaches – decade-pooled α and individualized α – and examined their effects on age-dependent metabolite profiles relative to commonly used approaches (the CSF-only and metabolite-pooled α). Applying the decade-pooled α and individualized α tissue correction approaches modified the metabolite-age relationships for mI, tCho and tNAA, and revealed nonlinear relationships between brain metabolism (mI and tCho) and age. In an additional analysis of potential sex-based differences, the decade-pooled α and individualized α correction approaches revealed that the increases in tCho and tCr with age may potentially be driven by female participants.

### Metabolite α values

Here, it was found that the α values for Glx, mI and tCr were below one, indicating higher concentrations in GM compared to WM, while the α values for tCho were greater than one, indicating greater levels of tCho in GM vs. WM, consistent with prior reports^52^. The α values for tNAA were around or slightly greater than one, indicating approximately equal levels in GM and WM, similar to what has been previously reported^10^. The α values for Glx, mI, tCho, and tCr were also found to differ significantly between different decades of life, suggesting that the concentration ratio of these metabolites in WM relative to GM changes with age. In contrast, tNAA α did not differ significantly between different decades of age, suggesting that the relative distribution across gray and white matter remains stable across the adult lifespan. Together, these findings indicate that the degree to which α changes with age is metabolite-specific, and that a single-grouped α fails to capture the wide range of individual α values present for each metabolite and decade of life.

### Metabolite levels across the lifespan

To date, the vast majority of ^1^H-MRS aging studies, including a previous study using the same dataset, have characterized age-dependent effects using linear models alone^26^. The present study addresses a major limitation of this approach by modeling metabolite cross-sectional “trajectories” as a nonlinear function of age which allows for the identification of timepoints of critical metabolic shifts across the lifespan. Such nonlinear shifts in neurobiology have been demonstrated with other neuroimaging modalities (e.g., fMRI and DTI)^27^ as well as select ^1^H-MRS studies of GABA across the adult lifespan^29^ and Glx, GABA+, tCr, tNAA, tCho, and mI from early childhood to early adulthood^65^. Our study contributes novel evidence of nonlinear metabolite shifts, i.e., nonlinear increases in mI, tCho, and tCr in females across the adult lifespan. While the biological basis for these nonlinear aging effects is not fully understood, hormonal transitions such as menopause could contribute as these transitions represent periods of heightened vulnerability in brain structure, connectivity, and energy metabolism^11^. Additional biological explanations for the non-linear aging effects and other metabolite-age relationships as a function of tissue correction approach are elaborated on below:

#### Glx

The Glx signal reflects glutamate, the brain’s primary excitatory neurotransmitter and contributor to cellular energy metabolism, and glutamine, which astrocytes produce from excess glutamate to prevent excitotoxicity^66,67^. Consistent with some prior reports in the PCC^26^ and hippocampus^68^, no significant age effects were found for Glx. In contrast, other studies have reported significant declines in Glx with age, including in the dorsal medial prefrontal cortex^28^ and calcarine cortex^69^. Despite these mixed findings, the overall stability of Glx with age observed here may reflect homeostatic regulation. Astrocytes clear and metabolize excess synaptic glutamate for energy, helping maintain stable Glx levels despite age-related changes in glutamatergic activity^70^. It should be noted that although no significant age effects were found for Glx, the slope of the linear age-related decline in Glx seen with the CSF-only correction was significantly reduced once α tissue correction approaches were applied. As such, some of decline in Glx seen with the CSF-only correction may reflect a partial-volume effect driven by age-related GM atrophy instead of true metabolic change.

#### mI

mI is considered an astroglial marker, with higher concentrations in astrocytes than neurons^71^. Because mI helps regulate astrocyte volume, its levels rise with astrocytes activation, making elevated mI a commonly used marker of neuroinflammation^72^. Prior ^1^H-MRS studies using CSF-only correction have reported mixed findings with some studies reporting no significant change in mI with age across several cortical regions, including the dorsal medial prefrontal cortex^28^, anterior cingulate cortex^73^, and PCC^73^. In contrast, other studies have reported significant mI increases with age^68,74^. In this study, nonlinear increases in mI with age were found with both the decade-pooled α correction and individualized α correction but not the CSF-only or metabolite-pooled α correction, although this finding was statistically significant only for the decade-pooled α correction. This accelerated increase in mI at mid-life may reflect increasing glial growth and reactive astrogliosis^75^. This is consistent with previous reports linking higher mI during midlife to greater cardiovascular dementia risk burden, driven partly by increasing age and amyloid accumulation, both of which may promote glial activation^76^.

#### tCho

The signal of tCho combines phosphocholine (PCh), a precursor in membrane phospholipid and myelin synthesis^77^, and glycerophosphocholine (GPC), a breakdown product phospholipid catabolism^78^, and is thus considered a marker of overall membrane turnover. Consistent with prior literature, tCho was found to increase with age across all tissue correction methods^26,28,73,79^, which may reflect greater membrane turnover and neuroinflammation with increasing age^80–82^. It should be noted, however, that the decade-pooled α and individualized α corrections revealed that this tCho-age relationship is nonlinear, which was not observable with the age-dependent slopes derived from the CSF-only or metabolite-pooled α corrected data. Upon closer inspection of potential sex-based differences, this nonlinear increase in tCho with age appear to be driven primarily by female participants. This female-specific pattern may reflect declining estrogen levels around menopause, which reduces estrogen’s protective effect on membrane integrity. One prior study, for example, found that postmenopausal women not on estrogen replacement therapy (ERT) show higher Cho than both long-term ERT users and younger women^83^. Further analysis of individualized α corrected concentrations across age groups revealed higher tCho in younger males than in younger females, and lower tCho in younger females than in older females, consistent with increases in tCho being largely driven by estrogen loss in older female participants.

#### tCr

The ^1^H-MRS tCr signal originates from both creatine and phosphocreatine, which are responsible for regenerating ATP and buffering energy supply in the brain^84^. The overall tCr increase with age across all methods is consistent with previous reports and may reflect increased glial proliferation with age, since glial cells have higher creatine and choline than neurons^71,85–87^, as well as compensatory shifts in energy metabolism due to creatine’s role in ATP buffering^69^. Upon further examination of individualized α corrected tCr concentrations across age groups, we observed that tCr was higher in older female participants than in younger female participants but not in older male participants vs. younger male participants. Thus, these increases in tCr with age may be primarily driven by female participants and related to estrogen’s role in brain energy metabolism. One study in perimenopausal women, for instance, has found that lower serum estradiol was associated with higher brain creatine, specifically in gray matter, suggesting that declining estradiol may cause a compensatory increase in creatine as an energy supply^88^.

#### tNAA

The tNAA signal is a combination of both NAA, a marker of neuronal integrity^89^, and NAAG, a modulator of glutamate release that may contribute to neuroprotection and synaptic plasticity^90^. Here, declines in tNAA with age were found when using a CSF-only, metabolite-pooled α, and decade-pooled α correction approaches but not the individualized α correction, and appeared to be driven primarily by male participants in the former three approaches. Nonetheless, this age-related decline is consistent with prior work and may be driven primarily by decreases in NAA due to declining neuronal and mitochondrial health with increasing age^24,28,91–94^. This male-specific decline may be due to sex differences that have been observed in mitochondrial function. Previous studies have reported greater mitochondrial respiratory capacity in females compared to males, including higher citrate synthase activity, a maker of mitochondrial content, and lower production of reactive oxygen species^95–98^. Since NAA is synthesized in neuronal mitochondria, these differences may help explain why we observed lower tNAA in males, as having less protection against oxidative stress could lead to a greater age-related decline in neuronal integrity.

### Tissue correction approach

Taken together, our findings indicate that the choice of tissue correction approach can significantly alter metabolite-age relationships. As such, the choice of tissue correction method is an important factor and must be selected carefully depending on the design of ^1^H-MRS study. Applying an overly specific tissue correction based on inaccurate or rigid assumptions can introduce noise and potentially remove interindividual or disease-related differences between groups. In longitudinal ^1^H-MRS studies for instance, the primary goal is to track metabolite changes in each individual using their own initial measurements as a baseline. If such a study population encompasses an epoch of the lifespan during which changes in bulk tissue composition are expected to be minimal (e.g., younger individuals), a CSF-only correction that preserves individual differences between timepoints may be the most appropriate. Correcting with a decade-pooled α, however, may be a more appropriate choice for case-control studies in cohorts where significant changes in bulk tissue composition are expected (e.g. neurodegenerative disorders) as this approach can account for partial volume effects without running the risk of over- or under-correcting with an individualized α or singular metabolite α which could mask or overestimate disease-related effects.

In the present study, the metabolite-age profiles with the CSF-only and metabolite-pooled α corrections were altered significantly when a decade-pooled α or individualized α correction was applied instead. As such, it is likely that both the CSF-only and metabolite-pooled α corrections are insufficient in correcting for bulk tissue changes in cross-sectional aging studies, such as the present study. In such cases, an individualized α tissue correction method may be more appropriate as this tailored approach could remove some inter-individual differences and mitigate some of the cohort effects that this type of study design is prone to. It should be noted, however, that accurately measuring individual α correction values requires a linear regression fit across metabolite levels measured from multiple regions in the same individual. Given the constraints of time-limited, costly scan sessions, only a limited number of regions may be sampled with multiple single-voxel acquisitions, unless a multi-voxel acquisition (e.g., MRSI) is performed. If an insufficient number of samples is acquired per person, some error may be introduced into the individualized α measurements and hence, the metabolite measurements. The present study estimated individualized α values from only two regions-of-interest per person which may have resulted in some error in the α value estimation and some of the non-significant relationships between metabolites and age as seen here (e.g. tNAA). Thus, in cases where MRS data cannot be acquired from multiple regions for each participant, a decade-pooled α correction may be more appropriate approach.

## Conclusion

In summary, four different tissue correction strategies were compared in a large healthy adult cohort with ages spanning five decades of life. The metabolite-age profiles for mI, tNAA, Glx, and tCho were the most sensitive to the choice of tissue correction method while the age-dependent profile for tCr was the least sensitive. The decade-pooled α and individualized α approaches revealed possible non-linear increases in mI and tCho with age that were not apparent when using the CSF-only or metabolite-pooled α correction approaches. Taken together, these findings indicate that the choice of tissue correction approach is an important factor to consider with designing cross-sectional aging studies as it can significantly alter metabolite-age relationships. The choice of tissue correction method should also be tailored to the specific study design and population, taking into consideration whether the study is cross-sectional or longitudinal and whether bulk tissues changes are expected in the cohorts-of-interest.

## Acknowledgments

We thank Richard Edden and Steve Hui for providing demographic information for the data used in this study.

## AI Statement

Generative AI (Claude, Anthropic) was used to help in writing R code implementing the Generalized Additive Models (GAMs).

## Ethics Statement

We analyzed de-identified neuroimaging data obtained from Shandong for which volunteers were recruited with local IRB approval.

## References

1. Benedetti, B. et al. Influence of aging on brain gray and white matter changes assessed by conventional, MT, and DT MRI. Neurology 66, 535–539 (2006).

2. Lv, L., Guo, H., Zhao, Z. C Zhao, X. Structural and microstructural changes in white and gray matter across the Alzheimer’s disease continuum. Front. Aging Neurosci. Volume 17-2025, (2025).

3. Lie, I. A. et al. Relationship Between White Matter Lesions and Gray Matter Atrophy in Multiple Sclerosis. Neurology 98, e1562–e1573 (2022).

4. Casella, C., Lipp, I., Rosser, A., Jones, D. K. C Metzler-Baddeley, C. A Critical Review of White Matter Changes in Huntington’s Disease. Mov. Disord. 35, 1302–1311 (2020).

5. Harris, A. D., Puts, N. A. J. C Edden, R. A. E. Tissue correction for GABA-edited MRS: Considerations of voxel composition, tissue segmentation, and tissue relaxations. J. Magn. Reson. Imaging 42, 1431– 1440 (2015).

6. Gasparovic, C. et al. Use of tissue water as a concentration reference for proton spectroscopic imaging. Magn. Reson. Med. 55, 1219–1226 (2006).

7. Gussew, A., Erdtel, M., Hiepe, P., Rzanny, R. C Reichenbach, J. R. Absolute quantitation of brain metabolites with respect to heterogeneous tissue compositions in 1H-MR spectroscopic volumes. Magn. Reson. Mater. Phys. Biol. Med. 25, 321–333 (2012).

8. Porges, E. C. et al. Impact of tissue correction strategy on GABA-edited MRS findings. NeuroImage 162, 249–256 (2017).

9. Pouwels, P. J. W. C Frahm, J. Regional metabolite concentrations in human brain as determined by quantitative localized proton MRS. Magn. Reson. Med. 39, 53–60 (1998).

10. Pouwels, P. J. W. C Frahm, J. Differential distribution of NAA and NAAG in human brain as determined by quantitative localized proton MRS. NMR Biomed. 10, 73–78 (1997).

11. Mosconi, L. et al. Menopause impacts human brain structure, connectivity, energy metabolism, and amyloid-beta deposition. Sci. Rep. 11, 10867 (2021).

12. Hädel, S., Wirth, C., Rapp, M., Gallinat, J. C Schubert, F. Effects of age and sex on the concentrations of glutamate and glutamine in the human brain. J. Magn. Reson. Imaging 38, 1480–1487 (2013).

13. Sailasuta, N., Ernst, T. C Chang, L. Regional variations and the effects of age and gender on glutamate concentrations in the human brain. Magn. Reson. Imaging 26, 667–675 (2008).

14. Vipin, A. et al. APOE4 carrier status determines association between white matter disease and grey matter atrophy in early-stage dementia. Alzheimers Res. Ther. 15, 103 (2023).

15. Klosinski, L. P., Yao, J., Yin, F., Fonteh, A. N., Harrington, M. G., Christensen, T. A., Trushina, E., C Brinton, R. D. White Matter Lipids as a Ketogenic Fuel Supply in Aging Female Brain: Implications for Alzheimer’s Disease - ClinicalKey. https://www.clinicalkey.com/#!/content/playContent/1-s2.0-S2352396415301924?returnurl=https:%2F%2Flinkinghub.elsevier.com%2Fretrieve%2Fpii%2FS2352396415301924%3Fshowall%3DtrueCreferrer=#hl0001243 (2015).

16. Human Gray and White Matter Metabolomics to Differentiate APOE and Stage Dependent Changes in Alzheimer’s Disease. J. Cell. Immunol. 3, (2021).

17. Yu, Y., Herman, P., Rothman, D. L., Agarwal, D. C Hyder, F. Evaluating the gray and white matter energy budgets of human brain function. J Cereb Blood Flow Metab 38, 1339–1353 (2018).

18. Maes, C. et al. Age-related differences in GABA levels are driven by bulk tissue changes. Hum. Brain Mapp. 39, 3652–3662 (2018).

19. Chen, X., Sachdev, P. S., Wen, W. C Anstey, K. J. Sex differences in regional gray matter in healthy individuals aged 44–48 years: A voxel-based morphometric study. NeuroImage 36, 691–699 (2007).

20. Kanchibhotla, S. C. et al. Genetics of ageing-related changes in brain white matter integrity – A review. Spec. Issue Invertebr. Models Aging 12, 391–401 (2013).

21. Parsaei, M., Sanjari Moghaddam, H. C Aarabi, M. H. Sex differences in brain structures throughout the lifetime. Aging Brain 4, 100098 (2023).

22. Martens, L. et al. Met carriers of the BDNF Val66Met polymorphism show reduced Glx/NAA in the pregenual ACC in two independent cohorts. Sci. Rep. 11, 6742 (2021).

23. Veldic, M. et al. Genetic variant in SLC1A2 is associated with elevated anterior cingulate cortex glutamate and lifetime history of rapid cycling. Transl. Psychiatry 9, 149 (2019).

24. Brooks, J. C. W. et al. A Proton Magnetic Resonance Spectroscopy Study of Age-related Changes in Frontal Lobe Metabolite Concentrations. Cereb. Cortex 11, 598–605 (2001).

25. Saunders, D. E., Howe, F. A., van den Boogaart, A., Griffiths, J. R. C Brown, M. M. Aging of the adult human brain: In vivo quantitation of metabolite content with proton magnetic resonance spectroscopy. J. Magn. Reson. Imaging 9, 711–716 (1999).

26. Gong, T. et al. Neurometabolic timecourse of healthy aging. NeuroImage 264, 119740 (2022).

27. Antal, B. B. et al. Brain aging shows nonlinear transitions, suggesting a midlife “critical window” for metabolic intervention. Proc. Natl. Acad. Sci. 122, e2416433122 (2025).

28. Ortuño, M. et al. The effect of age and sex on brain metabolites: from childhood to adulthood. NeuroImage 317, 121353 (2025).

29. Porges, E. C., Jensen, G., Foster, B., Edden, R. A. C Puts, N. A. The trajectory of cortical GABA across the lifespan, an individual participant data meta-analysis of edited MRS studies. eLife 10, e62575 (2021).

30. Endres, D. et al. On the Effect of Sex on Prefrontal and Cerebellar Neurometabolites in Healthy Adults: An MRS Study. Front. Hum. Neurosci. Volume 10-2016, (2016).

31. Hui, S. C. N. et al. The macromolecular MR spectrum does not change with healthy aging. Magn. Reson. Med. 87, 1711–1719 (2022).

32. Tkáč, I., Starčuk, Z., Choi, I.-Y. C Gruetter, R. In vivo 1H NMR spectroscopy of rat brain at 1 ms echo time. Magn. Reson. Med. 41, 649–656 (1999).

33. A Haase, J Frahm, W Hanicke, C D Matthaei. 1H NMR chemical shift selective (CHESS) imaging. Phys. Med. Biol. 30, 341 (1985).

34. Lin, A., et al. Minimum Reporting Standards for in vivo Magnetic Resonance Spectroscopy (MRSinMRS): Experts’ consensus recommendations. NMR Biomed. 34, e4484 (2021).

35. Lin, A., et al. Minimum Reporting Standards for in vivo Magnetic Resonance Spectroscopy (MRSinMRS): Experts’ consensus recommendations. NMR Biomed. 34, e4484 (2021).

36. Edden, R. A. E., Puts, N. A. J., Harris, A. D., Barker, P. B. C Evans, C. J. Gannet: A batch-processing tool for the quantitative analysis of gamma-aminobutyric acid–edited MR spectroscopy spectra. J. Magn. Reson. Imaging 40, 1445–1452 (2014).

37. Barkhuijsen, H., de Beer, R. C van Ormondt, D. Improved algorithm for noniterative time-domain model fitting to exponentially damped magnetic resonance signals. J. Magn. Reson. 1ScS 73, 553–557 (1987).

38. Borbath, T., Murali-Manohar, S., Dorst, J., Wright, A. M. C Henning, A. ProFit-1D—A 1D fitting software and open-source validation data sets. Magn. Reson. Med. 86, 2910–2929 (2021).

39. Simpson, R., Devenyi, G. A., Jezzard, P., Hennessy, T. J. C Near, J. Advanced processing and simulation of MRS data using the FID appliance (FID-A)—An open source, MATLAB-based toolkit. Magn. Reson. Med. 77, 23–33 (2017).

40. Govindaraju, V., Young, K. C Maudsley, A. A. Proton NMR chemical shifts and coupling constants for brain metabolites. NMR Biomed. 13, 129–153 (2000).

41. Govind, V., Young, K. C Maudsley, A. A. Corrigendum: Proton NMR chemical shifts and coupling constants for brain metabolites. Govindaraju V, Young K, Maudsley AA, NMR Biomed. 2000; 13: 129–153. NMR Biomed. 28, 923–924 (2015).

42. Wellcome Centre for Human Neuroimaging. Statistical Parametric Mapping (SPM12). University College London.

43. Friston, K. J. et al. Statistical parametric maps in functional imaging: A general linear approach. Hum. Brain Mapp. 2, 189–210 (1994).

44. Wansapura, J. P., Holland, S. K., Dunn, R. S. C Ball Jr., W. S. NMR relaxation times in the human brain at 3.0 tesla. J. Magn. Reson. Imaging 9, 531–538 (1999).

45. Lu, H. et al. Routine clinical brain MRI sequences for use at 3.0 Tesla. J. Magn. Reson. Imaging 22, 13– 22 (2005).

46. Piechnik, S. K., Evans, J., Bary, L. H., Wise, R. G. C Jezzard, P. Functional changes in CSF volume estimated using measurement of water T2 relaxation. Magn. Reson. Med. 61, 579–586 (2009).

47. Li, Y. et al. T1 and T2 Metabolite Relaxation Times in Normal Brain at 3T and 7T. *J*. Mol. Imaging 2, 1–5 (2013).

48. Träber, F., Block, W., Lamerichs, R., Gieseke, J. C Schild, H. H. 1H metabolite relaxation times at 3.0 tesla: Measurements of T1 and T2 values in normal brain and determination of regional differences in transverse relaxation. J. Magn. Reson. Imaging 19, 537–545 (2004).

49. Mlynárik, V., Gruber, S. C Moser, E. Proton T 1 and T 2 relaxation times of human brain metabolites at 3 Tesla. NMR Biomed. 14, 325–331 (2001).

50. Ernst, T., Kreis, R. C Ross, B. D. Absolute Quantitation of Water and Metabolites in the Human Brain. I. Compartments and Water. J. Magn. Reson. B 102, 1–8 (1993).

51. Leen, W. G., Willemsen, M. A., Wevers, R. A. C Verbeek, M. M. Cerebrospinal Fluid Glucose and Lactate: Age-Specific Reference Values and Implications for Clinical Practice. PLOS ONE 7, e42745 (2012).

52. Pouwels, P. J. W. C Frahm, J. Regional metabolite concentrations in human brain as determined by quantitative localized proton MRS. Magn. Reson. Med. 39, 53–60 (1998).

53. Huber, P. J. Robust Statistics. in International Encyclopedia of Statistical Science (ed. Lovric, M.) 1248–1251 (Springer Berlin Heidelberg, Berlin, Heidelberg, 2011). doi:10.1007/978-3-642-04898-2_594.

54. Wood, S. N. Fast Stable Restricted Maximum Likelihood and Marginal Likelihood Estimation of Semiparametric Generalized Linear Models. J. R. Stat. Soc. Ser. B Stat. Methodol. 73, 3–36 (2011).

55. Wood, S. N. Generalized Additive Models: An Introduction with R, Second Edition. (Chapman and Hall/CRC, 2017).

56. Wood, S. N., Pya, N. C Säfken, B. Smoothing Parameter and Model Selection for General Smooth Models. J. Am. Stat. Assoc. 111, 1548–1563 (2016).

57. Wood, S. N. Thin Plate Regression Splines. J. R. Stat. Soc. Ser. B Stat. Methodol. 65, 95–114 (2003).

58. Wood, S. N. Stable and Efficient Multiple Smoothing Parameter Estimation for Generalized Additive Models. J. Am. Stat. Assoc. 99, 673–686 (2004).

59. Hastie, T. C Tibshirani, R. Generalized Additive Models. Stat. Sci. 1, 297–310 (1986).

60. Feldman, D. et al. Flexible nonlinear modeling reveals age-related differences in resting-state functional brain connectivity in autistic males from childhood to mid-adulthood. Mol. Autism 16, 24 (2025).

61. Wilcoxon, F. Individual Comparisons by Ranking Methods. in Breakthroughs in Statistics: Methodology and Distribution (eds Kotz, S. C Johnson, N. L.) 196–202 (Springer New York, New York, NY, 1992). doi:10.1007/978-1-4612-4380-9_16.

62. Coretta, S. Tidygam: tidy prediction and plotting of generalized additive models. (2024).

63. Benjamini, Y. C Hochberg, Y. Controlling the False Discovery Rate: A Practical and Powerful Approach to Multiple Testing. J. R. Stat. Soc. Ser. B Methodol. 57, 289–300 (1995).

64. Moss, H. G., Jenkins, D. D., Yazdani, M. C Brown, T. R. Identifying the translational complexity of magnetic resonance spectroscopy in neonates and infants. NMR Biomed. 32, e4089 (2019).

65. Thomson, A. R. et al. The developmental trajectory of 1H-MRS brain metabolites from childhood to adulthood. Cereb. Cortex 34, bhae046 (2024).

66. Hubbard, J. A. C Binder, D. K. Chapter 9 - Glutamate Metabolism. in Astrocytes and Epilepsy 197–224 (Academic Press, San Diego, 2016). doi:10.1016/B978-0-12-802401-0.00009-0.

67. Andersen, J. V. C Schousboe, A. Milestone Review: Metabolic dynamics of glutamate and GABA mediated neurotransmission — The essential roles of astrocytes. J. Neurochem. 166, 109–137 (2023).

68. Yang, Z.-Y. et al. A quantitative analysis of 1H-MR spectroscopy at 3.0 T of three brain regions from childhood to middle age. Br. J. Radiol. 88, 20140693 (2015).

69. Salsano, I. et al. Neurochemical changes in GABA+, Glx, and the excitatory/inhibitory ratio in the calcarine cortex with healthy aging. NeuroImage 324, 121635 (2025).

70. Andersen, J. V. The Glutamate/GABA-Glutamine Cycle: Insights, Updates, and Advances. J. Neurochem. 169, e70029 (2025).

71. Brand, A., Richter-Landsberg, C. C Leibfritz, D. Multinuclear NMR Studies on the Energy Metabolism of Glial and Neuronal Cells. Dev. Neurosci. 15, 289–298 (1994).

72. Harris, J. L., Choi, I.-Y. C Brooks, W. M. Probing astrocyte metabolism in vivo: proton magnetic resonance spectroscopy in the injured and aging brain. Front. Aging Neurosci. Volume 7-2015, (2015).

73. Chiu, P.-W. et al. Metabolic changes in the anterior and posterior cingulate cortices of the normal aging brain: proton magnetic resonance spectroscopy study at 3 T. AGE 36, 251–264 (2014).

74. Chang, L., Ernst, T., Poland, R. E. C Jenden, D. J. In vivo proton magnetic resonance spectroscopy of the normal aging human brain. Life Sci. 58, 2049–2056 (1996).

75. Suri, S. et al. Effect of age and the APOE gene on metabolite concentrations in the posterior cingulate cortex. NeuroImage 152, 509–516 (2017).

76. Dounavi, M.-E. et al. Investigating the brain’s neurochemical profile at midlife in relation to dementia risk factors. Brain Commun. 6, fcae138 (2024).

77. Zhang, Y., Jen, F. E.-C., Fox, K. L., Edwards, J. L. C Jennings, M. P. The biosynthesis and role of phosphorylcholine in pathogenic and nonpathogenic bacteria. Trends Microbiol. 31, 692–706 (2023).

78. Tayebati, S. K. C Amenta, F. Choline-containing phospholipids: relevance to brain functional pathways. 51, 513–521 (2013).

79. Angelie, E. et al. Regional Differences and Metabolic Changes in Normal Aging of the Human Brain: Proton MR Spectroscopic Imaging Study. Am. J. Neuroradiol. 22, 119 (2001).

80. Duarte, J. M. N., Lei, H., Mlynárik, V. C Gruetter, R. The neurochemical profile quantified by in vivo 1H NMR spectroscopy. NEUROIMAGING THEN FUTURE 61, 342–362 (2012).

81. Chang, L., Ernst, T., Poland, R. E. C Jenden, D. J. In vivo proton magnetic resonance spectroscopy of the normal aging human brain. Life Sci. 58, 2049–2056 (1996).

82. Zahr, N. M., Mayer, D., Rohlfing, T., Sullivan, E. V. C Pfefferbaum, A. Imaging Neuroinflammation? A Perspective from MR Spectroscopy. Brain Pathol. 24, 654–664 (2014).

83. Robertson, D. M. W. et al. Effects of estrogen replacement therapy on human brain aging: An in vivo 1H MRS study. Neurology 57, 2114–2117 (2001).

84. Rackayova, V., Cudalbu, C., Pouwels, P. J. W. C Braissant, O. Creatine in the central nervous system: From magnetic resonance spectroscopy to creatine deficiencies. Introd. Vivo Magn. Reson. Spectrosc. MRS Method Non-Invasively Study Metab. 529, 144–157 (2017).

85. Mrak, R. E. C Griffin, W. S. T. Glia and their cytokines in progression of neurodegeneration. Dev. Orig. Aging Brain Blood Vessels 26, 349–354 (2005).

86. Finch, C. E. Neurons, glia, and plasticity in normal brain aging. Suppl. Brain Behav. Differ. Stages Hum. Life Proc. Expert Workshop Organ. Int. Health Found. Wassenaar Neth. 18-1S April 2002 24, S123–S127 (2003).

87. Urenjak, J., Williams, S., Gadian, D. C Noble, M. Proton nuclear magnetic resonance spectroscopy unambiguously identifies different neural cell types. J. Neurosci. 13, 981 (1993).

88. Ostojic, S. M. C Ostojic, J. Brain creatine, estradiol and neurocognitive complaints in perimenopausal women: an exploratory cross-sectional study. Neurosci. Lett. 880, 138616 (2026).

89. Castellano, G., Dias, C. S. B., Foerster, B., Li, L. M. C Covolan, R. J. M. NAA and NAAG variation in neuronal activation during visual stimulation. Braz. J. Med. Biol. Res. 45, (2012).

90. Coyle, J. T. The Nagging Question of the Function ofN-Acetylaspartylglutamate. Neurobiol. Dis. 4, 231–238 (1997).

91. Chiu, P.-W. et al. Metabolic changes in the anterior and posterior cingulate cortices of the normal aging brain: proton magnetic resonance spectroscopy study at 3 T. AGE 36, 251–264 (2014).

92. Cleeland, C., Pipingas, A., Scholey, A. C White, D. Neurochemical changes in the aging brain: A systematic review. Neurosci. Biobehav. Rev. 98, 306–319 (2019).

93. Schuff, N. et al. N-Acetylaspartate as a Marker of Neuronal Injury in Neurodegenerative Disease. in N-Acetylaspartate (eds Moffett, J. R., Tieman, S. B., Weinberger, D. R., Coyle, J. T. C Namboodiri, A. M. A.) 241–262 (Springer US, Boston, MA, 2006).

94. Mahmoudi, N. et al. Microstructural and Metabolic Changes in Normal Aging Human Brain Studied with Combined Whole-Brain MR Spectroscopic Imaging and Quantitative MR Imaging. Clin. Neuroradiol. 33, 993–1005 (2023).

95. Mishra, S. C Singh, K. K. Chapter 9 - Sex-specific differences in mitochondrial function and its role in health disparities. in Principles of Gender-Specific Medicine (Fourth Edition) (ed. Legato, M. J.) 129– 144 (Academic Press, 2023). doi:10.1016/B978-0-323-88534-8.00014-6.

96. Lagranha, C. J., Deschamps, A., Aponte, A., Steenbergen, C. C Murphy, E. Sex Differences in the Phosphorylation of Mitochondrial Proteins Result in Reduced Production of Reactive Oxygen Species and Cardioprotection in Females. Circ. Res. 106, 1681–1691 (2010).

97. Nagarajan-Radha, V., Aitkenhead, I., Clancy, D. J., Chown, S. L. C Dowling, D. K. Sex-specific effects of mitochondrial haplotype on metabolic rate in Drosophila melanogaster support predictions of the Mother’s Curse hypothesis. Philos. Trans. R. Soc. B Biol. Sci. 375, 20190178 (2019).

98. Silaidos, C. et al. Sex-associated differences in mitochondrial function in human peripheral blood mononuclear cells (PBMCs) and brain. Biol. Sex Differ. 9, 34 (2018).

