## Supplemental Table 1 for "Tissue correction strategies for ^1^H-MRS and their effects on age-dependent metabolite alterations"

**Supporting Information Table S1** Minimum Reporting Standards in Magnetic Resonance Spectroscopy checklist

Hardware

| | |
|---|---|
| Field Strength | 3T |
| Manufacturer | Philips Healthcare |
| Model (Software) | Ingenia CX |
| RF Coil | 32-channel head coil |
| Additional Hardware | Slice-selective lipid suppression band (20 mm thickness) |

Acquisition

| | |
|---|---|
| Pulse Sequence | PRESS |
| Volume of Interest (VOI) | PCC (posterior cingulate cortex) and CSO (centrum semiovale) |
| Nominal VOI size | 30 × 26 × 26 $mm^3$ (20.28 mL), same for both voxels |
| RF Coil | 32-channel head coil |
| Repetition Time (TR) and Echo Time (TE) | TR = 2 s and TE = 30 ms |
| Number of averaged spectra | 96 averages |
| Additional Parameters | 1.3 kHz refocusing bandwidth; 2000 Hz spectral width, 2048 spectral points |
| Water Suppression Method | VAPOR |
| Shimming Method | Pencil beam (PB)-auto, $2^{nd}$ order |
| Triggering or Motion Correction | N/A |

Analysis and Data Quality

| | |
|---|---|
| Analysis Software | Gannet 3.1 (preprocessing); Profit1D (spectral fitting) |
| Processing steps deviating from reference | HSVD residual water removal; zero-filling to 4096 points; 1 Hz exponential apodization; |

| | |
|---|---|
| | manual zero-order phase correction and frequency alignment when needed |
| Output Measure | Institutional Units |
| Quantification Reference | Unsuppressed water |
| Reported Variables | SNR (based off the fitted NAA peak); fit quality number (FQN, ratio of residual to noise variance) |
| Data exclusion criteria | Noisy spectra excluded through visual inspection (lack of interpretable spectrum) |
| Sample spectrum | Figures 1a and 1b |
